## Supplementary Information for "Molecular evidence of widespread benzimidazole drug resistance in *Ancylostoma caninum* from domestic dogs throughout the USA and discovery of a novel isotype-1 β-tubulin benzimidazole resistance mutation"

### S1 Appendix: *A. caninum* pooled samples across the USA.

The hookworm positive fecal samples collected from dogs across the USA were classified as "many", "moderate", "few", and "rare" based on IDEXX's semi-quantitative classification. Samples classified as "rare", as well as those classified as "few" but with <1g of feces available, were combined into pools before isolating the eggs. 65 such pooled samples, representing a total of 357 dogs across the USA, were sequenced at depth using the Illumina Miseq platform to enable the molecular detection of benzimidazole resistance from samples with low egg counts. Of these 65 pooled samples, the 293 bp and the 340 bp isotope-1 β-tubulin fragments encompassing codons 134, 167, 198, and 200 were successfully amplified from 63 samples. The samples were sequenced and analyzed using the same methods as described for the individual samples. The F167Y(TTC>TAC) and the novel Q134H(CAA>CAT) resistance mutations were present in 77.7% (49/63) and 53.9% (34/63) of the pooled samples, respectively. Their overall frequencies in the positive samples were 33.1% (95% C.I. 25.0% - 41.3%) and 10.2% (95% C.I. 5.3% - 15.1%), respectively (S6 Fig A,B). Variant calling at codons 198 and 200 revealed that 100% of the pooled samples contained susceptible alleles at these codons (data not shown).

### S1 Table: Information on pooled samples of *A. caninum*

The table contains information on the 65 pooled samples that were used in the study, their geographical region, the number of samples in each pool, and the number of eggs that were used for genomic DNA preparation.

| Pool | No. of Eggs | No. of Samples | Region |
|---|---|---|---|
| P1 | 1000 | 13 | NE |
| P10 | 800 | 2 | NE |
| P11 | 300 | 4 | NE |
| P12 | 500 | 8 | NE |
| P13 | 300 | 7 | NE |
| P14 | 1000 | 6 | NE |
| P15 | 1500 | 8 | NE |
| P16 | 1100 | 7 | NE |
| P17 | 300 | 3 | NE |
| P18 | 1200 | 2 | NE |
| P19 | 1100 | 6 | NE |
| P2 | 2000 | 21 | NE |
| P20 | 1600 | 2 | NE |
| P21 | 1600 | 3 | NE |
| P22 | 1800 | 7 | NE |
| P23 | 600 | 6 | NE |

| | | | |
|---|---|---|---|
| P25 | 1225 | 7 | NE |
| P26 | 2700 | 11 | NE |
| P27 | 850 | 2 | NE |
| P29 | 575 | 2 | NE |
| P3 | 375 | 6 | NE |
| P30 | 550 | 3 | MW |
| P31 | 600 | 2 | MW |
| P32 | 1200 | 6 | MW |
| P33 | 1800 | 9 | MW |
| P34 | 1650 | 8 | MW |
| P35 | 5000 | 8 | MW |
| P36 | 6800 | 5 | MW |
| P37 | 1000 | 5 | MW |
| P38 | 1925 | 5 | MW |
| P39 | 2275 | 9 | MW |
| P4 | 1000 | 10 | NE |
| P40 | 6775 | 2 | MW |
| P41 | 10350 | 2 | MW |
| P42 | 3900 | 5 | MW |
| P43 | 200 | NA | MW |
| P44 | 325 | 2 | MW |
| P45 | 1000 | 3 | S |
| P46 | 2000 | 4 | S |
| P47 | 450 | 3 | S |
| P48 | 475 | 3 | S |
| P49 | 525 | 6 | S |
| P5 | 1000 | 4 | NE |
| P50 | 400 | 6 | S |
| P51 | 7425 | 3 | S |
| P52 | 2350 | 2 | S |
| P53 | 500 | 2 | S |
| P54 | 1550 | 2 | S |
| P55 | 1650 | 2 | S |
| P56 | 8000 | 2 | S |
| P57 | 1250 | 2 | W |

| | | | |
|---|---|---|---|
| P58 | 2950 | 2 | MW |
| P59 | 300 | 2 | W |
| P6 | 600 | 9 | NE |
| P60 | 450 | 2 | MW |
| P62 | 550 | 3 | MW |
| P63 | 1450 | 3 | S |
| P64 | 800 | 2 | S |
| P66 | 3500 | 2 | S |
| P67 | 3100 | 3 | W |
| P68 | 3500 | 12 | S |
| P69 | 2000 | 12 | S |
| P7 | 800 | 6 | NE |
| P8 | 1000 | 2 | NE |
| P9 | 1000 | 6 | NE |

**S2 Table: Primers used for Sanger sequencing of the near full-length isotype-1 β-tubulin**

| Primer Name | Sequence |
|---|---|
| AC_BT1_San_152_F1 | GGTGAATCTGATCTGCAACTT |
| AC_BT1_San_173_R5 | CAAGTTGCAGATCAGATTCAC |
| AC_BT1_San_389_F2 | CTTGATGTAGTTCGCAAAGAGG |
| AC_BT1_San_409_R4 | CTCTTTGCGAACTACATCAAGG |
| AC_BT1_San_577_F3 | GTGGAGCCATACAATGCTAC |
| AC_BT1_San_964_R3 | TCCAAGCGAGTGAGTCAATTG |
| AC_BT1_San_1051_R1 | GTTCTTGTTCTGCACTGACATC |
| AC_BT1_San_2320_R2 | GTGTAGCATTGTATGGCTCCA |
| AC_BT1_San_2963_F4 | GATGTCAGTGCAGAACAAGA |

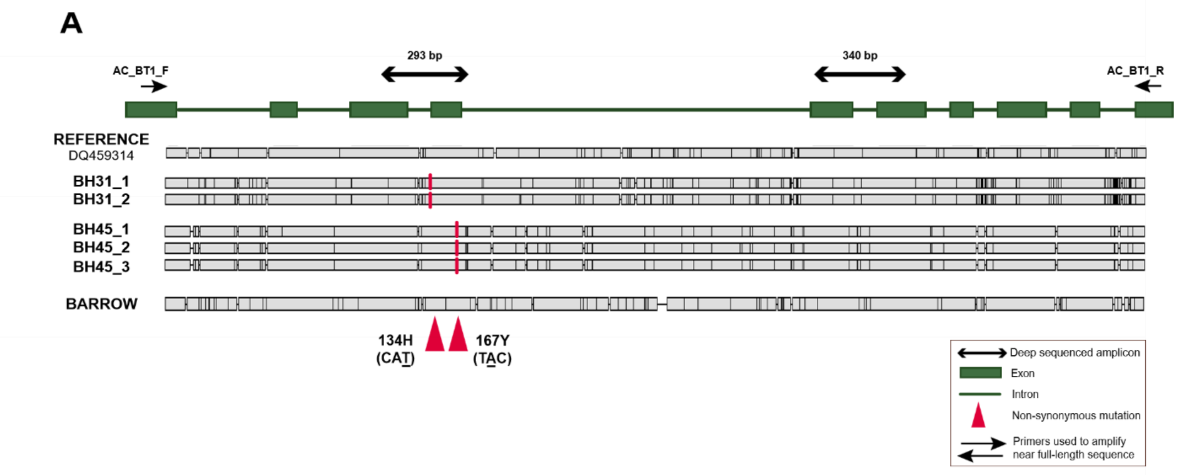


B

| | REFERENCE | BH31_1 | BH31_2 | BH45_1 | BH45_2 | BH45_3 | BARROW |
|---|---|---|---|---|---|---|---|
| REFERENCE | | 135 | 130 | 156 | 157 | 154 | 184 |
| BH31_1 | 135 | | **7** | 210 | 212 | 209 | 239 |
| BH31_2 | 130 | 7 | | 205 | 207 | 204 | 234 |
| BH45_1 | 156 | 210 | 205 | | **5** | **2** | 184 |
| BH45_2 | 157 | 212 | 207 | 5 | | **3** | 185 |
| BH45_3 | 154 | 209 | 204 | 2 | 3 | | 182 |
| BARROW | 184 | 239 | 234 | 184 | 185 | 182 | |

**S1 Fig: Full-length genomic sequences of the *A. caninum* isotype-1 β-tubulin gene showing the sequence polymorphisms present**

(A): Multiple Sequence Alignment of the 3,467 bp long *A. caninum* isotype-1 β-tubulin near full-length genomic sequence cloned and sequenced from greyhound isolates BH31 and BH45, the benzimidazole-susceptible lab isolate (Barrow), and the *A. caninum* isotype-1 β-tubulin reference sequence (Genbank accession: DQ459314). Following Sanger sequencing, the sequences of each clone were assembled *de novo* using the Geneious software and the assembled consensus sequences for each clone were aligned using the MUSCLE tool for multiple sequence alignment in Geneious v. 10.0.9. The synonymous and non-synonymous polymorphisms relative to the reference sequence, DQ459314, are highlighted in black and red, respectively.

(B): Distance matrix showing the pairwise number of nucleotide differences among the sequences.

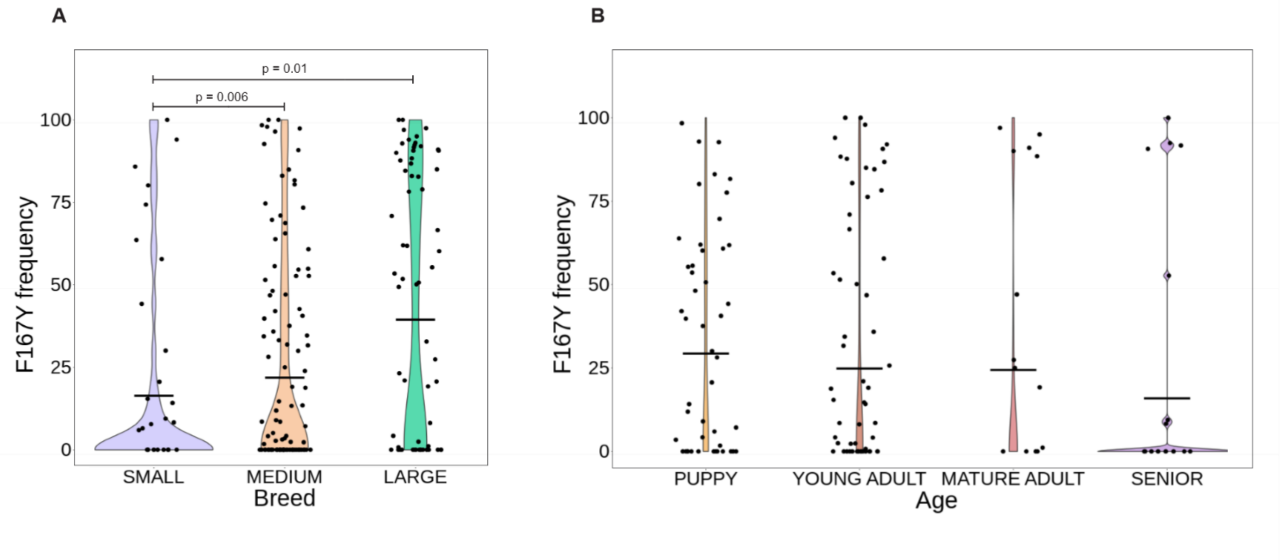


**S2 Fig: Frequency of the 167Y(T<u>A</u>C) resistance mutation in *A. caninum* from dogs of different breeds and age groups.**

(A) Violin Plot of the 167Y(T<u>A</u>C) resistance allele frequencies in *A. caninum* from dogs of different breed sizes. The mean frequency is indicated by a horizontal line and any statistically significant differences calculated using the pairwise Wilcoxon rank sum test ($p<0.05$) between the regions are indicated (p-value >0.05 not indicated).

(B) Violin Plot of the 167Y(T<u>A</u>C) resistance allele frequencies in *A. caninum* from dogs of different age groups. The mean frequency is indicated by a horizontal line and any statistically significant differences calculated using the pairwise Wilcoxon rank sum test ($p<0.05$) between the regions are indicated (p-value >0.05 not indicated).

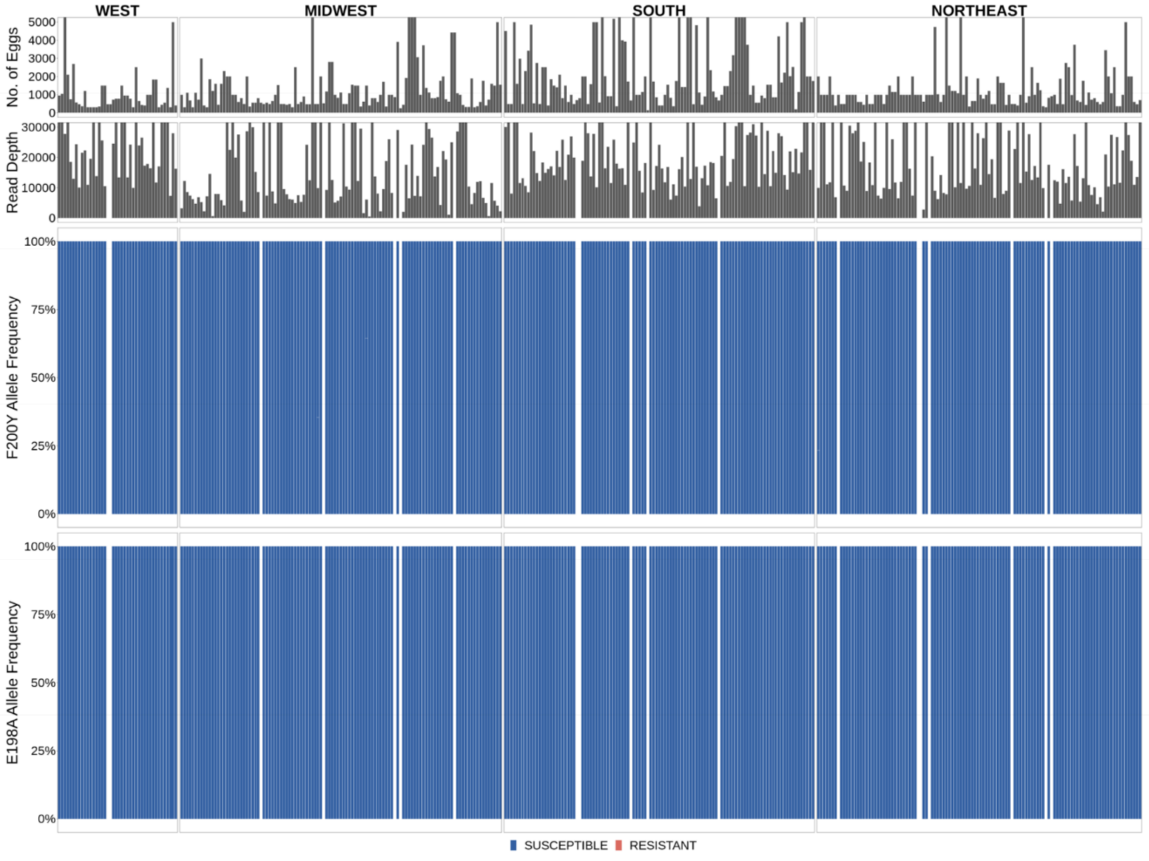


**S3 Fig: Absence of resistance mutations at codons 198 and 200 of the *A. caninum* isotype-1 β-tubulin gene**

Deep amplicon sequencing data of the 340 bp fragment of *A. caninum* from the 312/ 328 individual samples across the USA. The top chart is a histogram showing the number of eggs used to make each genomic DNA preparation, the middle chart is a histogram showing the mapped read depth for each sample, and the lower charts show the frequency of resistance or susceptible alleles present at codons 198 and 200 of the isotype-1 β-tubulin gene. Blue bars indicate the susceptible alleles 198E(GAA) and 200F(TTC) at both the codons.

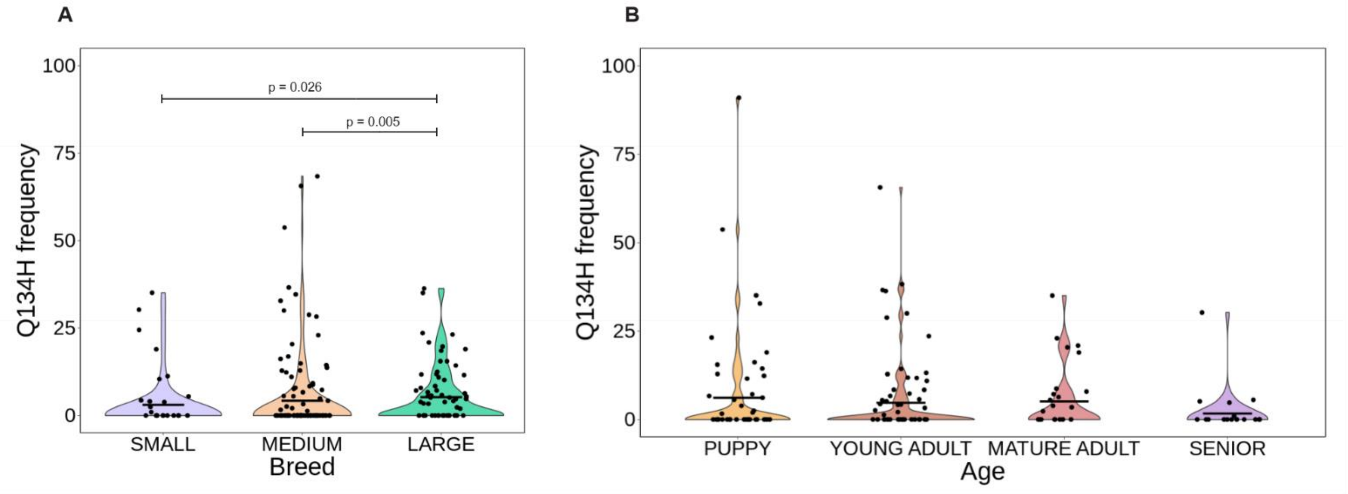


**S4 Fig: Frequency of the 134H(CAT) resistance mutation in *A. caninum* from dogs of different breeds and age groups.**

(A) Violin Plot of the 134H(CAT) resistance allele frequencies in *A. caninum* from dogs of different breed sizes. The mean frequency is indicated by a horizontal line and any statistically significant differences calculated using the pairwise Wilcoxon rank sum test ($p<0.05$) between the regions are indicated (p-value >0.05 not indicated).

(B) Violin Plot of the 134H(CAT) resistance allele frequencies in *A. caninum* from dogs of different age groups. The mean frequency is indicated by a horizontal line and any statistically significant differences calculated using the pairwise Wilcoxon rank sum test ($p<0.05$) between the regions are indicated (p-value >0.05 not indicated).

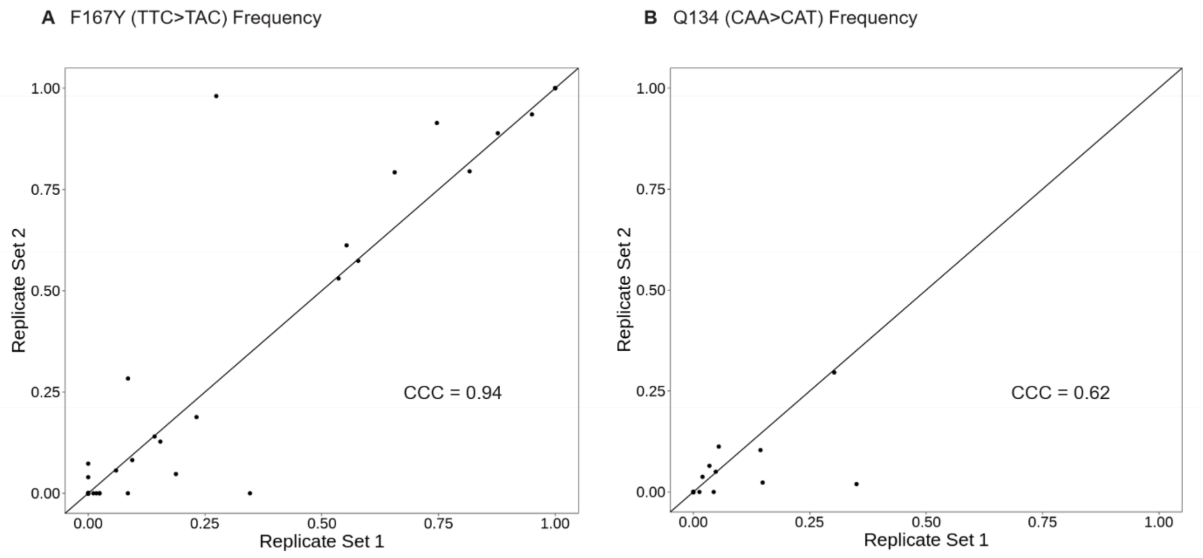


**S5 Fig: Assessment of the repeatability of SNP frequencies determined by amplicon sequencing**

(A): Scatterplot comparing the 167Y(TAC) resistance mutation frequencies determined from two independent replicate PCRs and Miseq runs for 50 samples. 24/50 samples carried the resistance mutation. The solid diagonal line indicates perfect agreement between the two runs. Lin’s Concordance Correlation Coefficient (CCC) indicates the level of agreement between the two sets of replicates.

(B): Scatterplot comparing the 134H(CAT) resistance mutation frequencies determined from two independent replicate PCRs and Miseq runs for 50 samples. 10/50 samples carried the resistance mutation. The solid diagonal line indicates perfect agreement between the two runs. Lin’s Concordance Correlation Coefficient (CCC) indicates the level of agreement between the two sets of replicates.

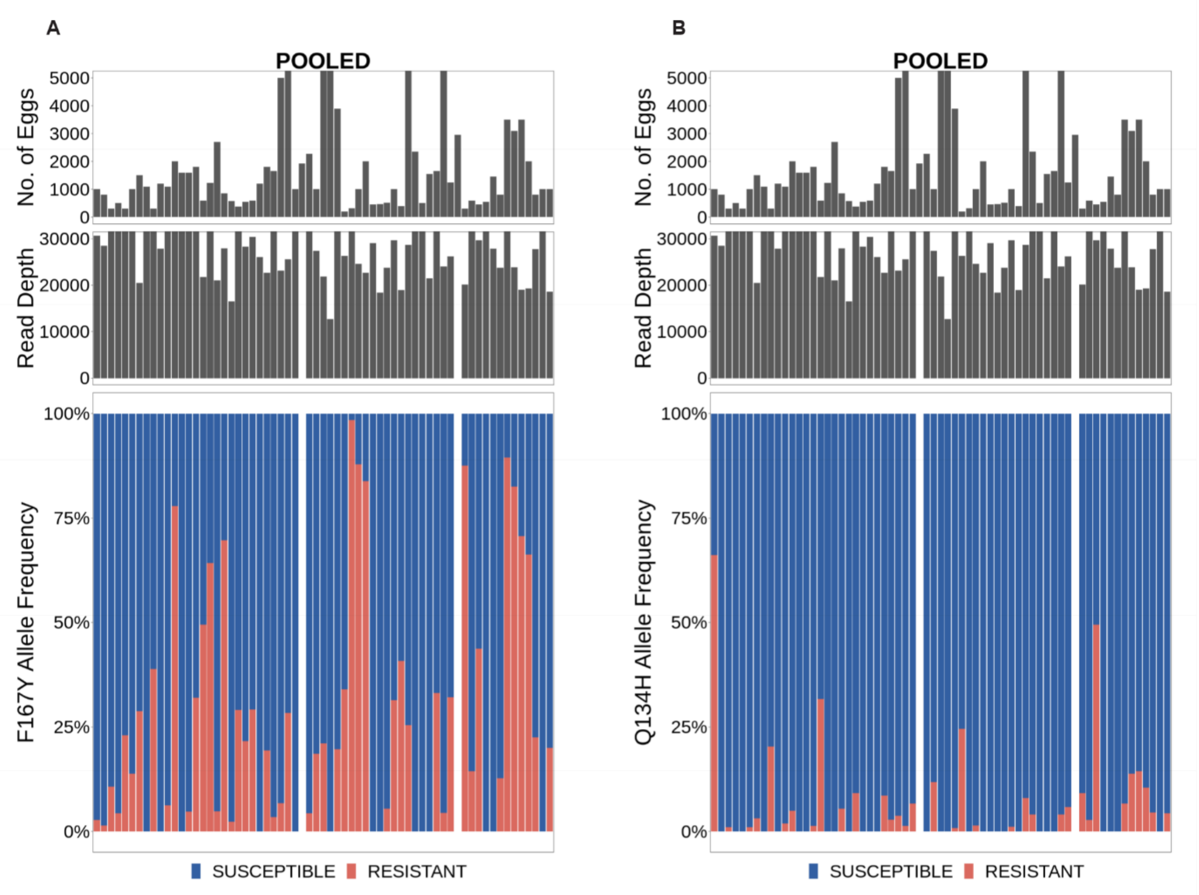


**S6 Fig: Frequency of the benzimidazole resistance mutations in pooled samples of *A. caninum* across the USA.**

(A): Deep amplicon sequencing data of the 293 bp isotype-1 β-tubulin fragment of *A. caninum* from the 63/ 65 pooled samples across the USA. The top chart is a histogram showing the number of eggs used to make each genomic DNA preparation, the middle chart is a histogram showing the mapped read depth for each sample, and the lower chart shows the relative frequency of the F167Y(TTC>TAC) mutation. Red bars indicate the 167Y(TAC) resistance allele and the blue bars indicate the 167F(TTC) susceptible allele.

(B): Deep amplicon sequencing data of the 293 bp isotype-1 β-tubulin fragment of *A. caninum* from the 63/ 65 pooled samples across the USA. The top chart is a histogram showing the number of eggs used to make each genomic DNA preparation, the middle chart is a histogram showing the mapped read depth for each sample, and the lower chart shows the relative frequency of the Q134H(CAA>CAT) mutation. Red bars indicate the 134H(CAT) resistance allele and the blue bars indicate the 134Q(CAA) susceptible allele.
